## Extended data figures for "Structural polymorphism of α-synuclein fibrils alters pathway of Hsc70 mediated disaggregation"

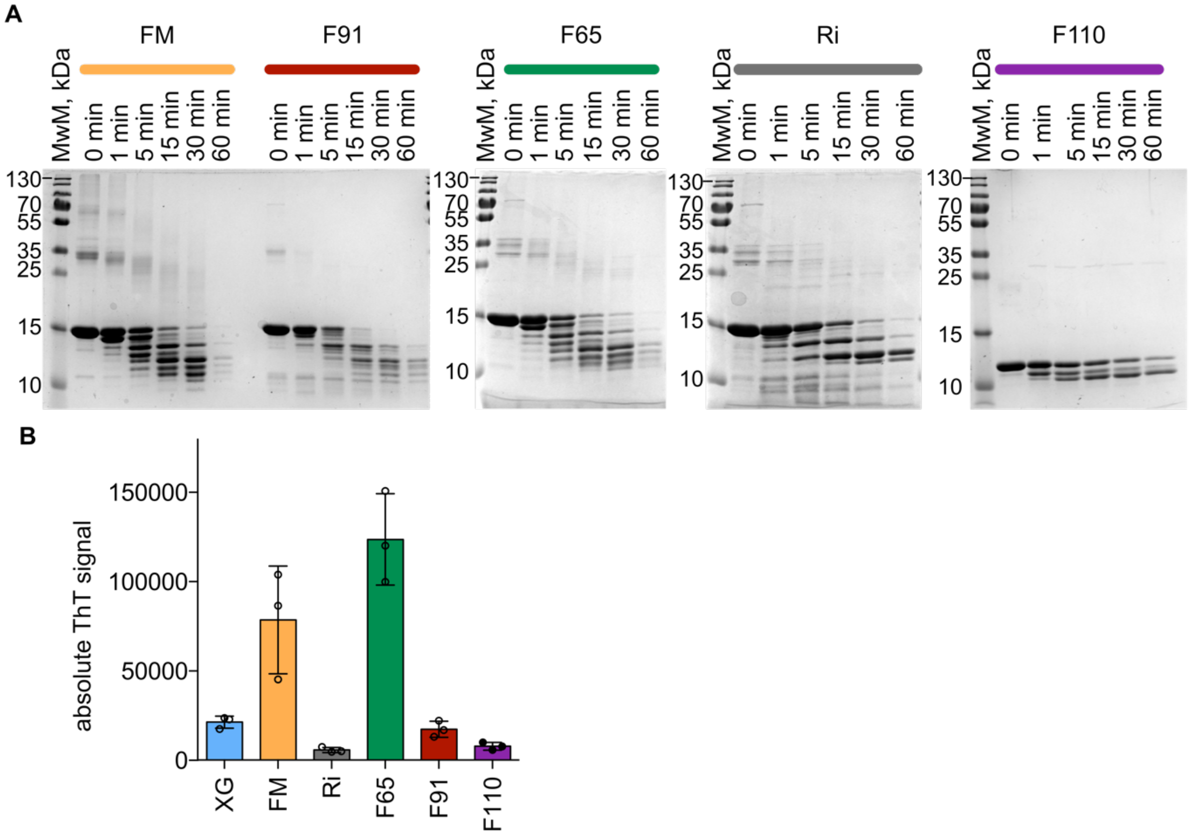


**Extended Data Figure 1 | $\alpha$-syn polymorphs differ in sensitivity to proteolysis and reactivity to Thioflavin T.**

**A** SDS-PAGE gels of limited proteolysis of FM (yellow), F91 (red), F65 (green), Ri (grey), and the C-terminally truncated F110 (purple) by proteinase K at timepoint 0, and after 1, 5, 15, 30 and 60 min. **B** Absolute Thioflavin T (ThT) signal of $\alpha$-syn fibrillar polymorphs (XG (blue), FM (yellow), Ri (grey), F65 (green), F91 (red), and the C-terminally truncated F110 (purple)) at the same concentration (2 µM protein – 30 µM ThT). Data are mean ± s.e.m. of three biological replicates.

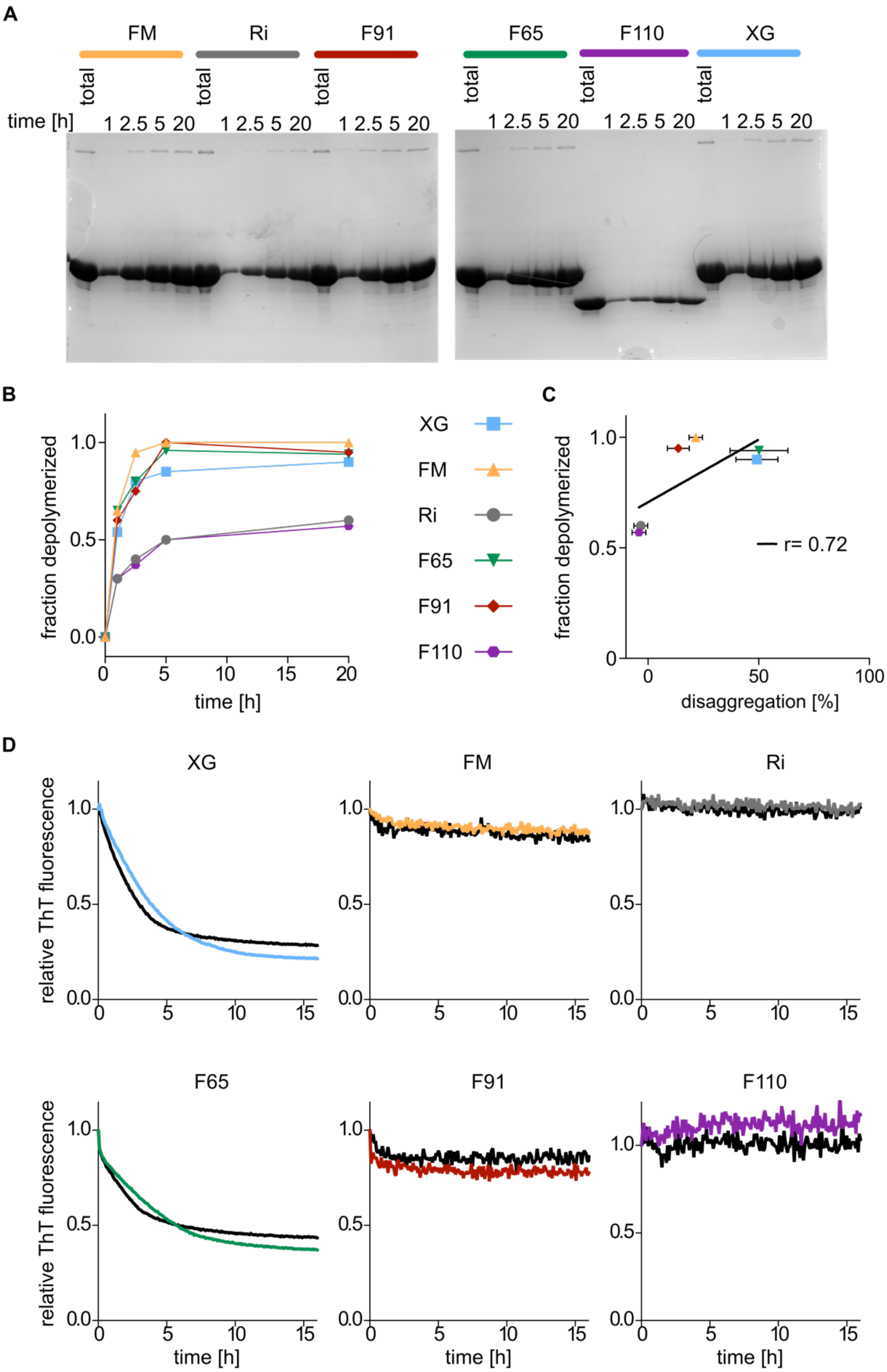


**Extended Data Figure 2 | High chemical stability of ribbons and F110 polymorphs correlates with their resistance to chaperone-mediated disaggregation**

**A** SDS-PAGE of $\alpha$-syn monomers released to supernatant, separated from fibrillar material by centrifugation, after incubation of fibrils on ice for the indicated times. **B** Fraction of total fibrils depolymerized, as shown in A, as a function of incubation time on ice. **C** Correlation plot of disaggregation after 16 hours in percent and fraction depolymerized with a linear correlation fit (solid line (r=0.72)). **D** ThT disaggregation assay with 1x (XG, blue; F91, red; F65, green; FM, yellow; Ri, grey; F110, purple) or 2x (black) chaperone concentration. Representative graphs of three technical replicates are shown.

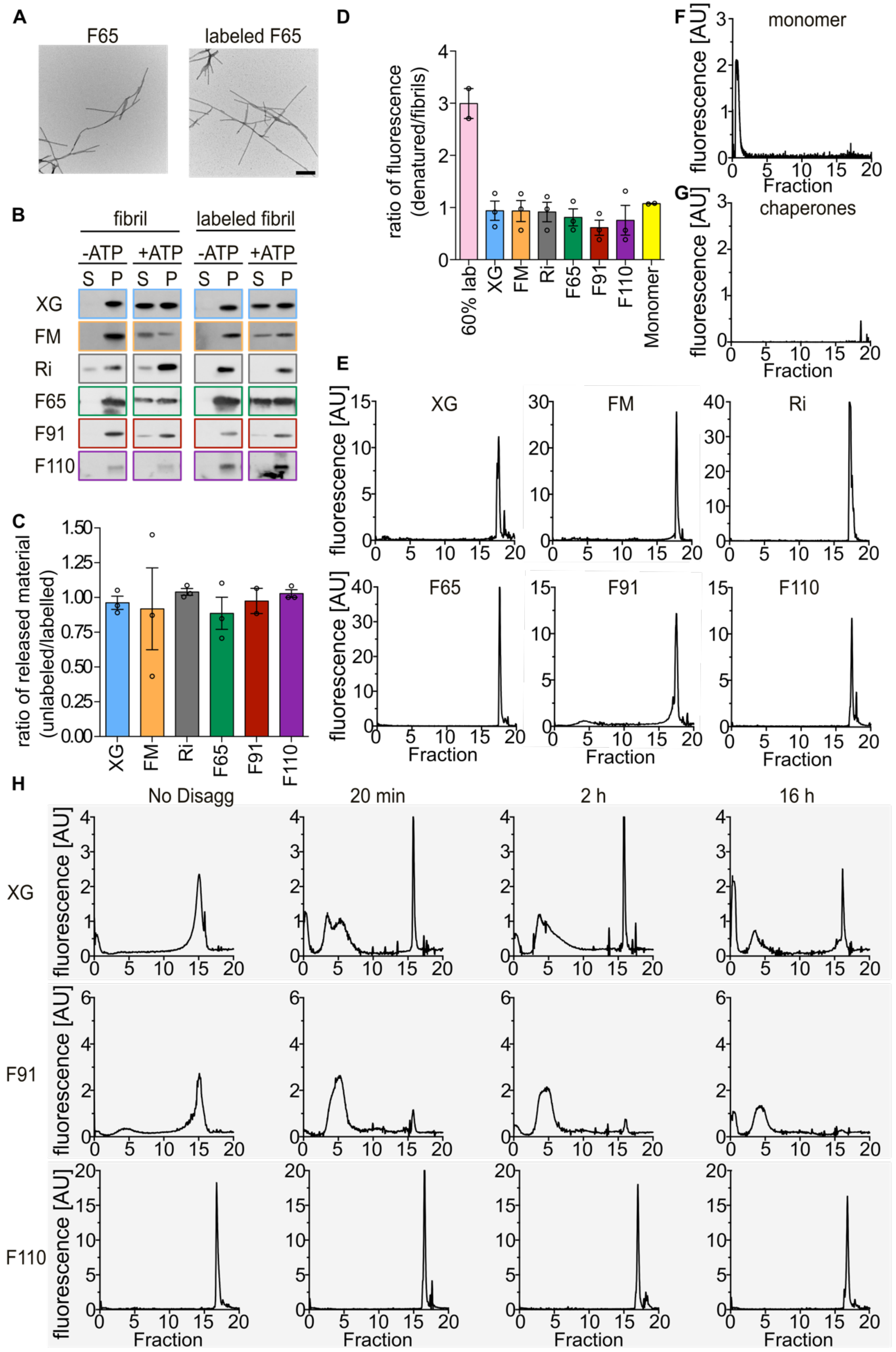


**Extended Data Figure 3 | Fibrillar fragments accumulate to various degrees during chaperone mediated disaggregation of polymorphs.**

**A** Negative stain EM images of polymorphs F65 unlabelled (left) and AF555-labeled (right) (scale bar 200 nm). **B** Representative western blot images of unlabelled (left) und AF555-labeled polymorphs (right) incubated with the chaperone machinery in the presence (+ATP) and absence of ATP (-ATP). **C** Ratio of released $\alpha$-syn protein of AF555-labeled and unlabelled fibrils of all polymorphs. Quantification of protein in the supernatant of the total (P + S) after 16h disaggregation by the active chaperone machinery in B (analysed with ImageJ). Data are mean ± s.e.m. **D** Ratio of normalized fluorescence of denatured AF555-labeled fibrils/monomer in GnHCl compared to labelled fibrils/monomer in buffer. **E** and **F** Sucrose density gradient (10-85%) profile of AF555-labeled monomers (**E**) and chaperones (Hsc70, DnaJB1, Apg2) only (**F**). Sucrose gradient of labelled fibrils (XG, FM, Ri, F65, F91, F110), are shown in **G**. **H** Representative sucrose gradient (10-85%) profile of a polymorph XG, F65, F91 and F110 after incubation of AF555-labelled fibrils with the active chaperone machinery (Hsc70, DnaJB1, Apg2, + ATP) for 20 min, 2 h, 16 h and 16 h incubation with the inactive machinery (-ATP, No Disaggregation).

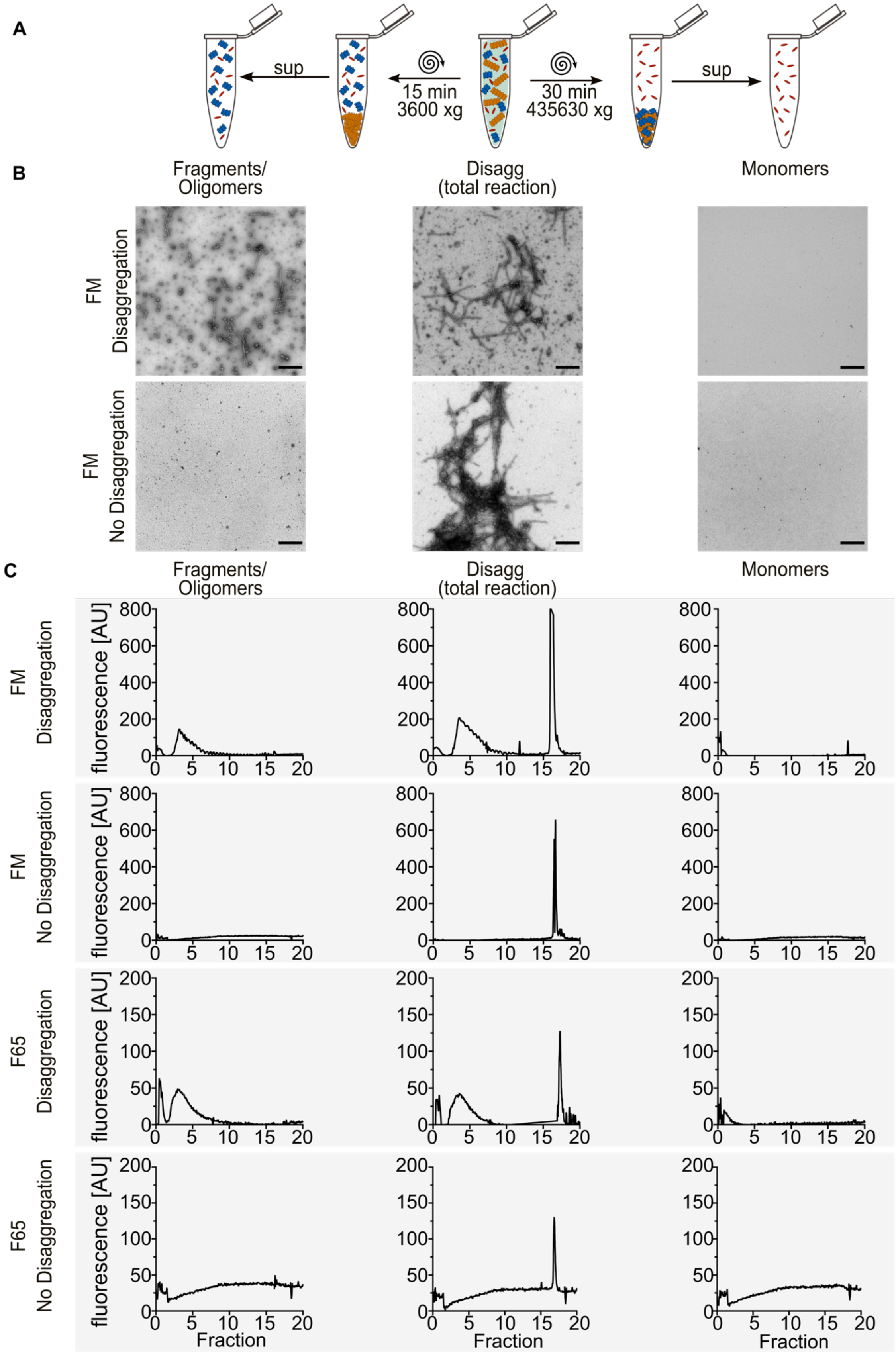


**Extended Data Figure 4 | Disaggregation reaction products can be fractioned based on differential centrifugation.**

**A** Centrifugation procedure as shown in Fig. 4C to separate different disaggregation reaction products. A total disaggregation reaction is centrifuged at 3,600g for 15 min, fibrils (orange) are separated in the pellet and small fragments/oligomers remain in the supernatant (left). By centrifugation of the total reaction at a higher speed (435,630g) for 30 min, small fragments/oligomers (blue) and fibrils are pelleted and only monomers (red) remain in the supernatant (right). **B** Representative electron micrographs of fibrils incubated with the chaperone machinery in the presence (Disaggregation) and absence (No Disaggregation) of ATP for the polymorph FM. Scale bar 500 nm. **C** Sucrose density gradient (10-85%) profile of AF555-labelled fibrils incubated with the active (Disaggregation) and inactive (No Disaggregation) chaperone machinery and similar prepared as in panel A.

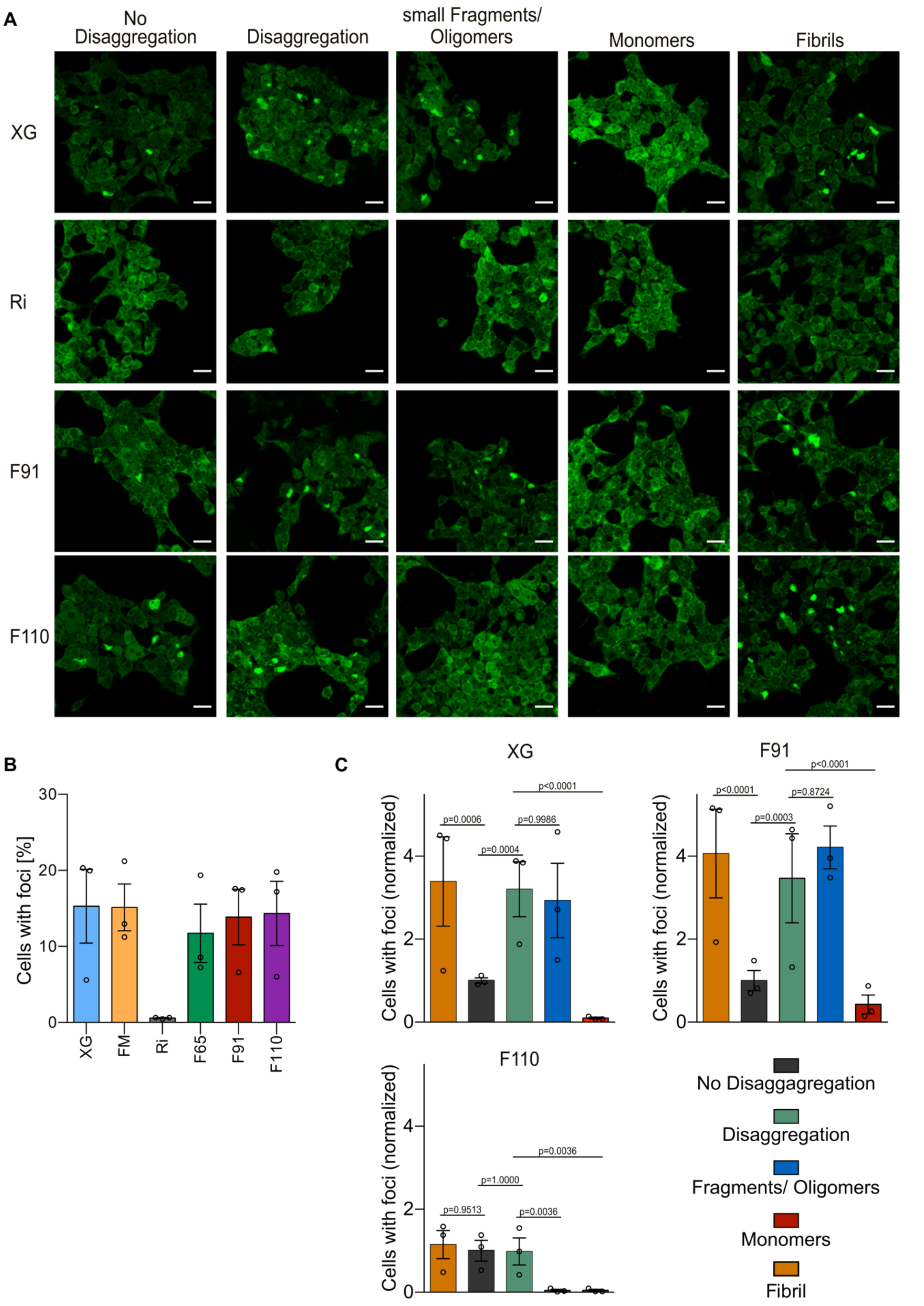


**Extended Data Figure 5| Disaggregation reaction products trigger foci formation in human cell model.**

**A** Representative fluorescence intensity microscopic images of HEK293T cells stably expressing $\alpha$-syn A53T-YFP seeded treated with fibril preparations of polymorph XG, Ri, F91 and F110. The cells were exposed to fibrils only, fibrils incubated with chaperones in the absence (No Disaggregation) and presence of ATP (Disaggregation), as well as small fragments/oligomer and monomer fractions of the disaggregation reaction separated by centrifugation as described in figure 4C. **B** Quantification of cells with foci normalized to the No Disaggregation sample (No disaggregation, black; Disaggregation, green; small fragments/oligomers, blue; monomers, red; fibrils, orange). The Ri polymorph did not induce aggregation in the used reporter cell system (in B) and was therefore not further analysed. Data are mean ± s.e.m. Statistical analysis was performed by nonparametric Two-Way ANOVA with pairwise comparisons of estimated marginal means with Tukey correction for multiple comparisons.

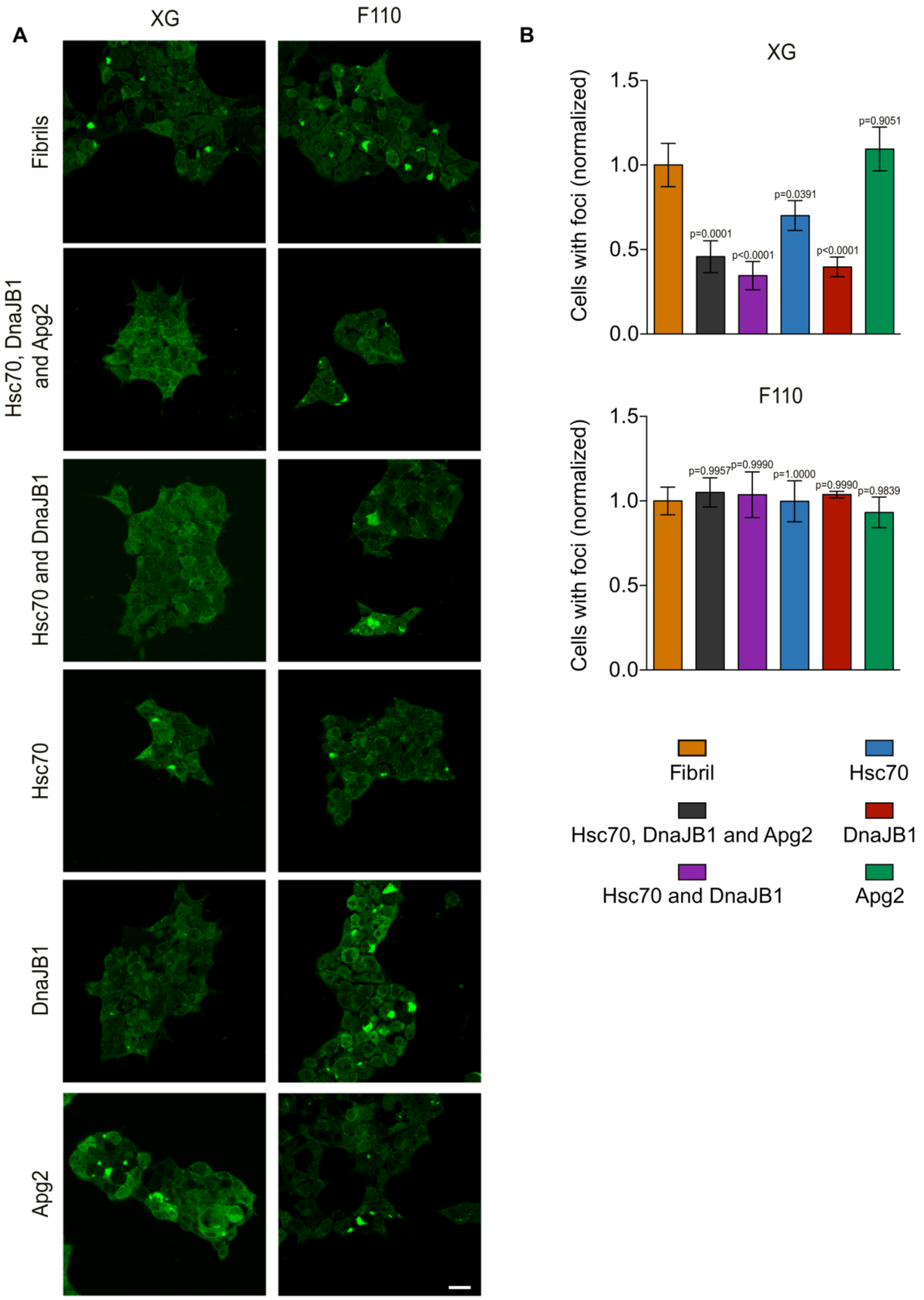


**Extended Data Figure 6| Pre-incubation of $\alpha$-syn fibrils with DNAJB1 reduces foci formation.**
**A** Representative fluorescence intensity microscopic images of HEK293T cells stably expressing $\alpha$-syn A53T-YFP seeded with fibrils (polymorphs XG and F110) incubated with different chaperone combinations; fibrils without chaperones as a control, fibrils with the whole chaperone machinery (Hsc70, DnaJB1, Apg2), Hsc70 and DnaJB1 and Hsc70, DnaJB1 and Apg2 individually (scale bar 20 µm).
**B** Quantification of normalized cell counts with foci (XG, top; F110, bottom). Data are mean ± s.e.m. Statistical analysis was performed by nonparametric Two-Way ANOVA with pairwise comparisons of estimated marginal means with Tukey correction for multiple comparisons.
